# Enhancing Statistical Power While Maintaining Small Sample Sizes in Behavioral Neuroscience Experiments Evaluating Success Rates

**DOI:** 10.1101/2024.07.25.605060

**Authors:** Theo Desachy, Marc Thevenet, Samuel Garcia, Anistasha Lightning, Anne Didier, Nathalie Mandairon, Nicola Kuczewski

## Abstract

Studies with low statistical power reduce the probability of detecting true effects and often lead to overestimated effect sizes, undermining the reproducibility of scientific results. While several free statistical software tools are available for calculating statistical power, they often do not account for the specialized aspects of experimental designs in behavioral studies that evaluate success rates. To address this gap, we developed “SuccessRatePower” a free and user-friendly power calculator based on Monte Carlo simulations that takes into account the particular parameters of these experimental designs. Using “SuccessRatePower”, we demonstrated that statistical power can be increased by modifying the experimental protocol in three ways:

1) reducing the probability of succeeding by chance (chance level), 2) increasing the number of trials used to calculate subject success rates, and, in some circumstance, 3) employing statistical analyses suited for discrete values. These adjustments enable even studies with small sample sizes to achieve high statistical power. Finally, we performed an associative behavioral task in mice, confirming the simulated statistical advantages of reducing chance levels and increasing the number of trials in such studies

## Introduction

Statistical power is the likelihood of obtaining a statistically significant result from a sample when the effect in the entire population equals or exceeds the minimum effect targeted by the study. In other words, it represents the probability that the p-value of a statistical test will be lower than the α-level when the alternative hypothesis is true (p-value < α/HA) (Neyman & Pearson, 1933; Cohen, 1962). Experimental studies with low statistical power have several negative implications. Firstly, they risk overlooking a genuine effect, leading to a false negative conclusion. Secondly, they compromise the precision of estimating population parameters from the experimental sample, potentially resulting in an overestimation of the detected effect when the statistical test is deemed significant (Button et al., 2013; Colquhoun, 2014; Curran-Everett, 2017). Thirdly, they diminish the positive predictive value, which refers to the proportion of published positive results that actually reflect a true effect (Ioannidis, 2005; Button et al., 2013; Colquhoun, 2014). Consequently, studies with low statistical power contribute to the ongoing reproducibility crisis in scientific research (Munafò et al., 2017; Ellis, 2022).

Statistical power is influenced by both the effect size (ES) in the studied populations and the sample size (Markel, 1991, Cohen, 1992; Cohen, 2002). While increasing the sample size is the most common method to enhance statistical power, it incurs additional costs in terms of money and time. Furthermore, larger sample sizes can also raise ethical concerns, particularly in invasive human and animal research studies (Sneddon et al., 2017; Andrade, 2020). However, adjustments to parameters other than sample size can also affect statistical power, depending on the study design (Brandmaier et al., 2015). For example, utilizing a repeated measurement design, where the modification of a variable within the same subject is followed overtime, can bolster statistical power (Vickers, 2003; Guo et al., 2013). Mathematical considerations suggest that studies replicating the measurement of the parameter of interest, with each subject being measured multiple times under each condition, may also enhance statistical power. However, this increase in power is contingent upon the absence of perfect correlations between the measurements, meaning that the data collected are not identical for all replications (Goulet and Cousineau, 2019).

Determining statistical power often demands a level of statistical expertise that may exceed the capabilities of many researchers (Fernandes-Taylor et al., 2011; Sullivan et al., 2016). Fortunately, several freely available software packages have been developed to facilitate power calculations (Faul et al., 2007; Lakens & Caldwell, 2021, NC3rs), with G*Power 3 being arguably the most comprehensive and widely utilized by the scientific community. However, these software tools can sometime overlook the peculiarities of specific research fields. One such scenario arises in behavioral studies, where differences in cognitive abilities (e.g., memory capabilities) are assessed by evaluating the subject success rate on a specific associative task (see, for example: Ravel et al., 1994; Devore & Linster, 2012; Alonso et al., 2012; Meier et al., 2016; Bratch et al., 2016; Mandairon et al., 2018; Grelat et al., 2018; Zhang et al., 2021).

A classical design for these types of studies is depicted in Figure 1. Here, each subject in the sample must perform several trials (replications) of a task, in which they must select the correct answer from multiple choices. Here the probability of succeeding at the task by chance, defined as the chance level, is determined by the ratio of correct choices to incorrect ones. In this type of protocol, the experimenter can vary three different parameters: 1) the number ‘n’ of subjects performing the task, 2) the number ‘t’ of trials each subject performs, and 3) the chance level. Furthermore, three different analytical procedures can be applied to the data. The first involves calculating a global population success rate by dividing the total number of successes by the total number of trials. Alternatively, one can first calculate the success rate of each subject and then determine the average success rate of the population. Finally, a statistical model for clustered data can be used to account for the nested nature of the data, i.e., the repetition of trials within the same subject. In the first case, a statistical test for discrete values (e.g., a proportion test) is used. In the second case, a statistical test for continuous values is required (e.g., a Student’s t-test). In the third case, multilevel models or generalized estimating equations can be applied. (Ballinger, 2004; Aarts *et al*., 2014; Olvera Astivia *et al*., 2019; but see McNabb & Murayama, 2021).

**Figure 1.**
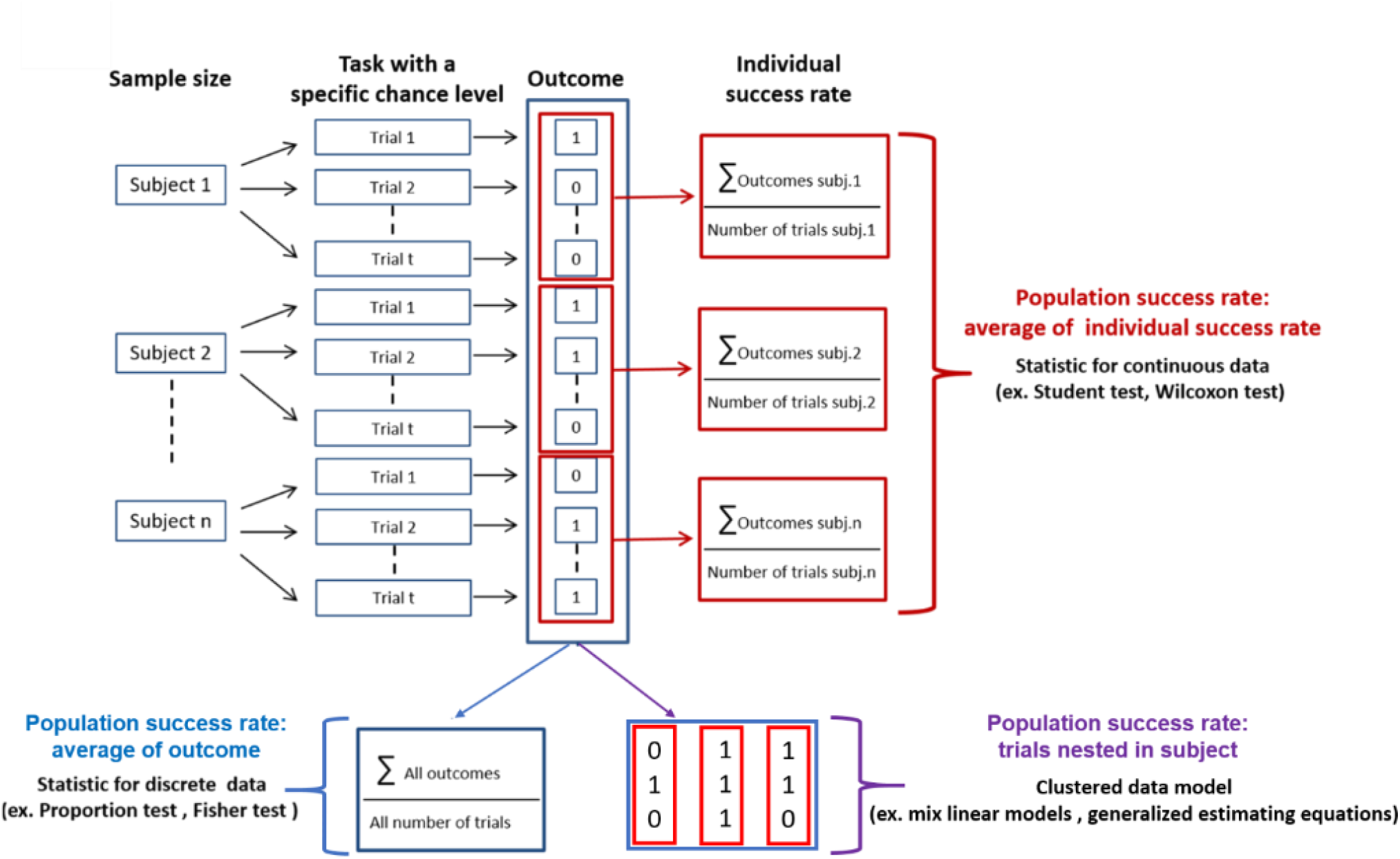
Typical design of experimental protocols aimed to determine the success rate for a behavioral task performed on *n* subjects and *t* trials (replication) per subject. The chance level (probability of making a correct choice) is equal to the number of correct choices divided by the total available choices. Outcome 0 represents failure, and outcome 1 represents success. Three statistical analytical procedures can be used to determine whether the success rate is different from chance: 1-Statistical tests on the individual success rates (shown in red). 2-Statistical tests on the population success rate (shown in blue). 3-Utilization of statistical models for clustered data that take into account the nested nature of the data (shown in purple). In all cases, the outcome for trained groups is compared to the chance level or to the success rate of a control group (not shown). Other than the modification of the sample size, the statistical power can potentially be affected by the modification of the chance level, the number of trials per subject, and the statistical analysis procedure adopted.

Importantly, the modification of the chance level is expected to change the statistical power only when the experimental group is compared to chance or when one of the two groups being compared performs at chance level. To clarify this point, consider the following scenario:

*We hypothesize that a specific brain region is necessary to support a certain form of associative memory. We predict that a targeted intervention, such as the removal of this brain region or a pharmacological treatment, will impair the retrieval of associative memory. To test this hypothesis, we design an experiment where both the control group and the treated group are trained on an associative memory task, resulting in an average success rate of 70% for both groups. After training, the intervention is applied to the treated group, and associative memory retrieval is subsequently tested in both groups*.

Three possible outcomes can arise from this experiment

### 1. The intervention prevents memory retrieval

In this scenario, the performance of the treated group will return to chance level. Since the effect size (ES) is the distance of the success rate between the control and treated groups its magnitude will therefore change as a function of the chosen chance level leading to the increase of statistical power (Figure 2, left).

**Figure 2:**
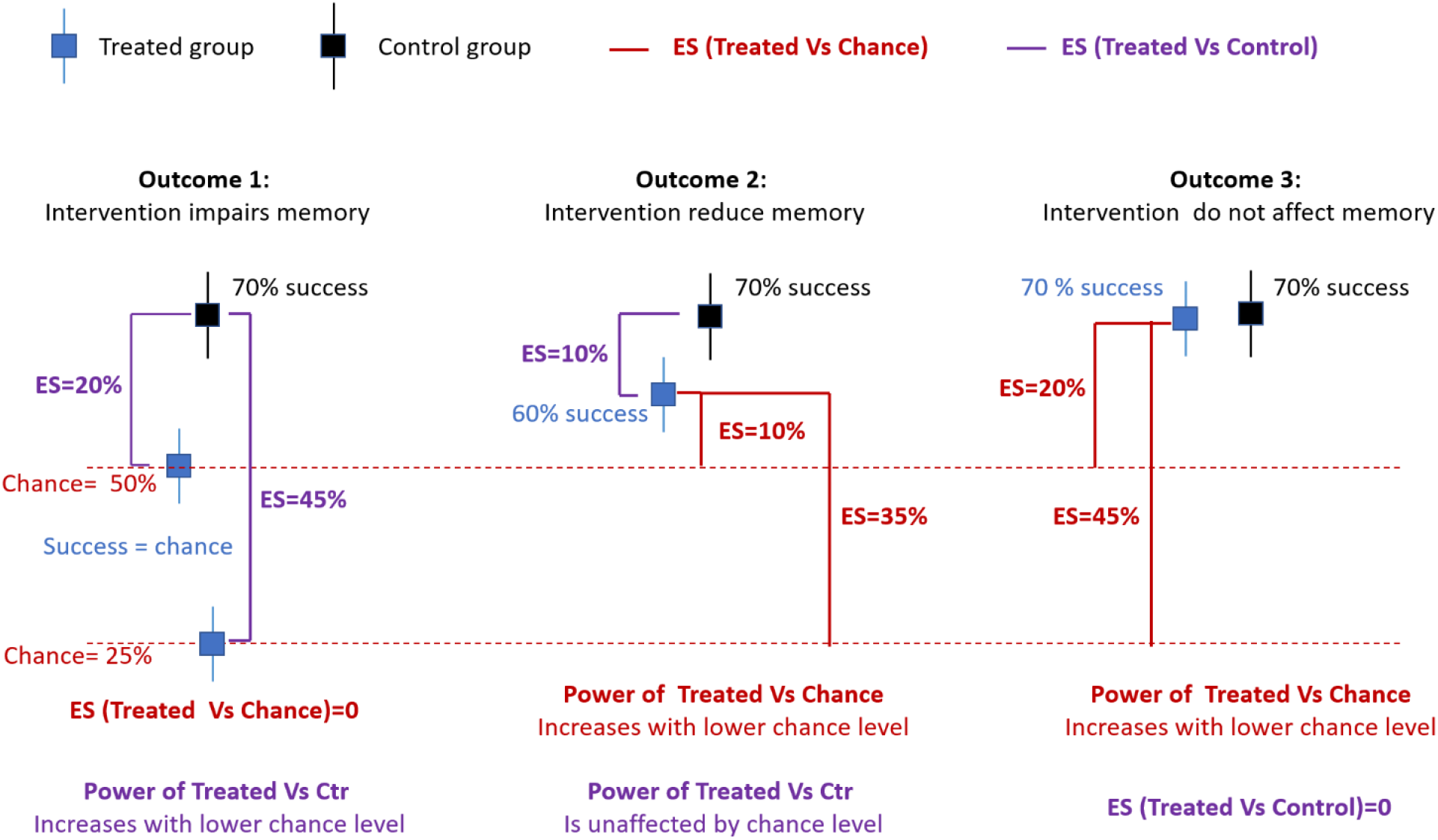
Impact of modifying the chance level on statistical power as a function of the outcome of the experiment assessing memory retrieval. ES = effect size

### 2. The intervention reduces but does not prevent memory retrieval

Here, the treated group will perform worse than the control group but better than chance level. In this condition the difference between the success rates of the two groups is now independent of the chosen chance level. However, it also important to assess whether the treated group perform better than chance and the statistical power of this comparison again depended on the chosen chance level (Figure 2, middle).

### 3. The intervention does not affect memory retrieval

In this scenario, the ES quantifying the difference between treaded and control group is zero and independent of chance level but again is important to verify that treated groups perform better than chance, and the statistical power of this compareson depend on the chose chance level (Figure 2, right).

The high flexibility inherent in this design type calls for a tool that enables researchers to compute and compare statistical power relative to the experimental strategies employed. As far as we are aware, such a tool has not yet been developed. Therefore, we present “SuccessRatePower” a freely accessible and user-friendly power calculator based on Monte Carlo simulation, tailored for these experimental protocols. We utilized SuccessRatePower to generate and compare statistical power across various experimental scenarios, identifying conditions that enhance statistical power. Subsequently, we implemented a behavioral protocol in mice to evaluate whether the animals’ performance was influenced by the application of experimental parameters anticipated to yield higher-powered designs.

## Material and methods

### Statistical simulation

Statistical power was calculated using Monte Carlo simulation. In the first step, the script randomly sampled the success probability for each subject in the sample. Since the success rate cannot exceed 100% or fall below the chance level, it generated random samples using the reduced Beta distribution. This approach allows for flexible modeling of continuous variables that are inherently bounded between a minimum and a maximum value, while also targeting specific population-level means and standard deviations. For that given user-specified values for the target population mean (μ), target population standard deviation (*σ*), and the bounds (minimum and maximum values), the script rescaled these parameters to the standard [0, 1] interval using the following transformations:

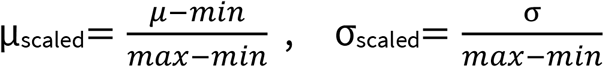

It then estimated the shape parameters α and β of the Beta distribution by minimizing the squared difference between the target and the theoretical mean and standard deviation of the Beta distribution. The optimal parameters were obtained using the scipy.optimize.minimize function in Python, with the constraint that both α and β must be positive. Once α and β were estimated, the script drew random samples from the Beta distribution using scipy.stats.beta.rvs(α, β, size=n), where n is the desired sample size. These values were then scaled back to the original range [min, max] as follows: X=min+Xbeta⋅(max−min). Then, the script used the sampled success probability of each subject in a binomial distribution to generate the outcome of the task (failure = 0, success = 1) for each of the subject’s trials. Finally, the outcomes of the two groups in each simulated sample were compared using the chosen statistical model. If the p-value of the comparison was lower than the predetermined α level, the result was considered a true positive when the average success rates of the populations from which the groups were drawn differed, and a false positive when they did not. This procedure was repeated 5,000 times. The simulation uses one-sided tests to assess the hypothesis that the trained group performs better than either the control group or the chance level. When the average probability of the trained group is set higher than the chance level, the script classifies any significant result (p-value < α) as a true positive. Conversely, when the average probability is the same for both groups, the script uses a two-sided test and significant results are classified as false positives. Statistical power and Type I error were estimated as the proportions of true positives and false positives across all repetitions, respectively:

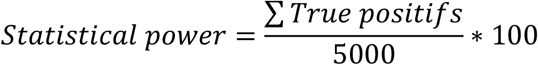

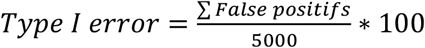

A flowchart illustrating the simulation process is provided in Supplementary Figure 1.The statistical simulation was conducted using the Python script available at the following link: (Script, executable SuccessRatePower).

The input parameters for the script are detailed in Table 1

**Table 1:**
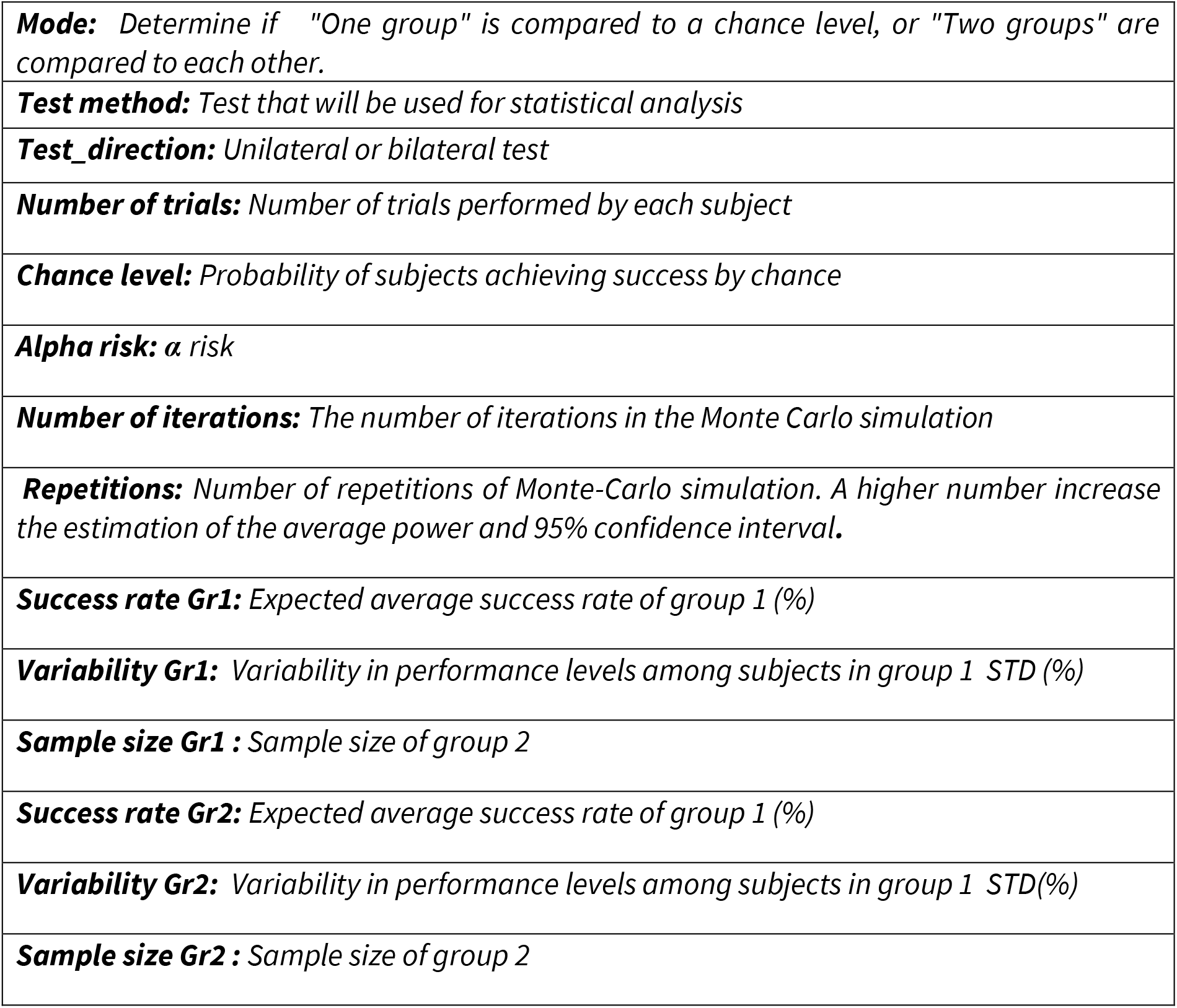
Input parameters of SuccessRatePower.

For the majority of the results presented later in this article, the average success probability of the trained group was set at 70% with 20% variability of performance between subjects. These values are compatible with those reported in the literature, where the success rate of healthy training groups falls between 70% and 90% (see Mandairon et al., 2018). For the analytical method based on continuous values, the success rate for each subject is calculated as the ratio of the number of successes to the number of trials. In this condition, we employed the SciPy function stats.ttest_1samp to compare the success rate of the trained group with the chance level. For comparing the sample success rate of the trained group with the success rate of the control group, we utilized the SciPy function stats.ttest_ind. For the analytical method based on discrete values, the script calculated the total number of success for the sample. In this condition, “statsmodels.stats.proportion.proportions_ztest” is used to compare the success rate of the trained group with both the chance level and success rate of the control group. Multilevel Mixed Model, were implemented using the Python function: smf.mixedlm(“success ~ group”, df, groups=df[“animal_id”], re_formula=“~1”); Generalized Estimating Equations (GEE), were implemented using the Python function: smf.gee(“success ~ group”, groups=“animal_id”, data=df, family=sm.families.Binomial())

#### Model Quality Check

The repeatability of the Monte Carlo simulation was assessed by repeating the power calculation 100 times with the same parameters and evaluating the standard deviation of the calculated power as a function of the number of iterations. As shown in Supplementary Figure 2, a variability lower than 1% was observed starting from 5,000 iterations.

To evaluate the capacity of SuccessRatePower to accurately estimate statistical power, we compared its performance to that of GPower 3 under the specific experimental and analytical conditions where power calculation is feasible with GPower 3—namely, a proportion z-test comparing two groups with no intersubject variability where the sample size is set as the number of subjects multiplied by the number of trials These comparisons, shown in Supplementary Figure 3, demonstrate similar performance between the two methods.

### Behavioral experiments (pre-registered protocols at osf)

#### Animals

Six adult male C57BL/6J mice (Janvier-Labs, France), aged 8 weeks at the beginning of the experiments, were housed in groups of six in standard laboratory cages with water ad libitum and maintained on a 12-hour light/dark cycle at a constant temperature of 22°C. Experiments were conducted following procedures in accordance with the European Community Council Directive of 22nd September 2010 (2010/63/UE) and adhered to the standards of the National Ethics Committee (Agreement APAFIS#29948-2021021812246456 v3). Animals were handled according to the ‘Mouse Handling: Tutorial’ provided by NC3Rs. Sample size was determine by an open-ended Sequential Bayes Factor design starting with 6 mice per group and add new subject until BF01<1/3 or BF01>3.

#### Learning and Experimental Design

The mice were trained to associate a food reward (a small amount of sweetened cereal from Kellogg’s, Battle Creek, MI, USA, of approximatively 1 cm thickness and 3–4 cm of diameter) with the presence of the odorant limonene+, within a radial maze containing either two or four arms (Figure 3). The odorant was introduced at the end of one of the maze arms through a custom-made olfactometer. Training sessions commenced after a shaping period lasting two days. During the shaping period, the animals were allowed to freely navigate the maze without any scent present and with the food reward randomly placed at the end of one of the arms. To incentivize the animals to complete the task, access to food in their home cage was limited to 2g of food pellets per mouse for the whole duration of the study, resulting in an approximately 20% reduction in daily consumption and a consequent 10% approximate reduction in body weight (Mandairon et al., 2018). The shaping period consist of two 5-minute exploration periods each day.

**Figure 3.**
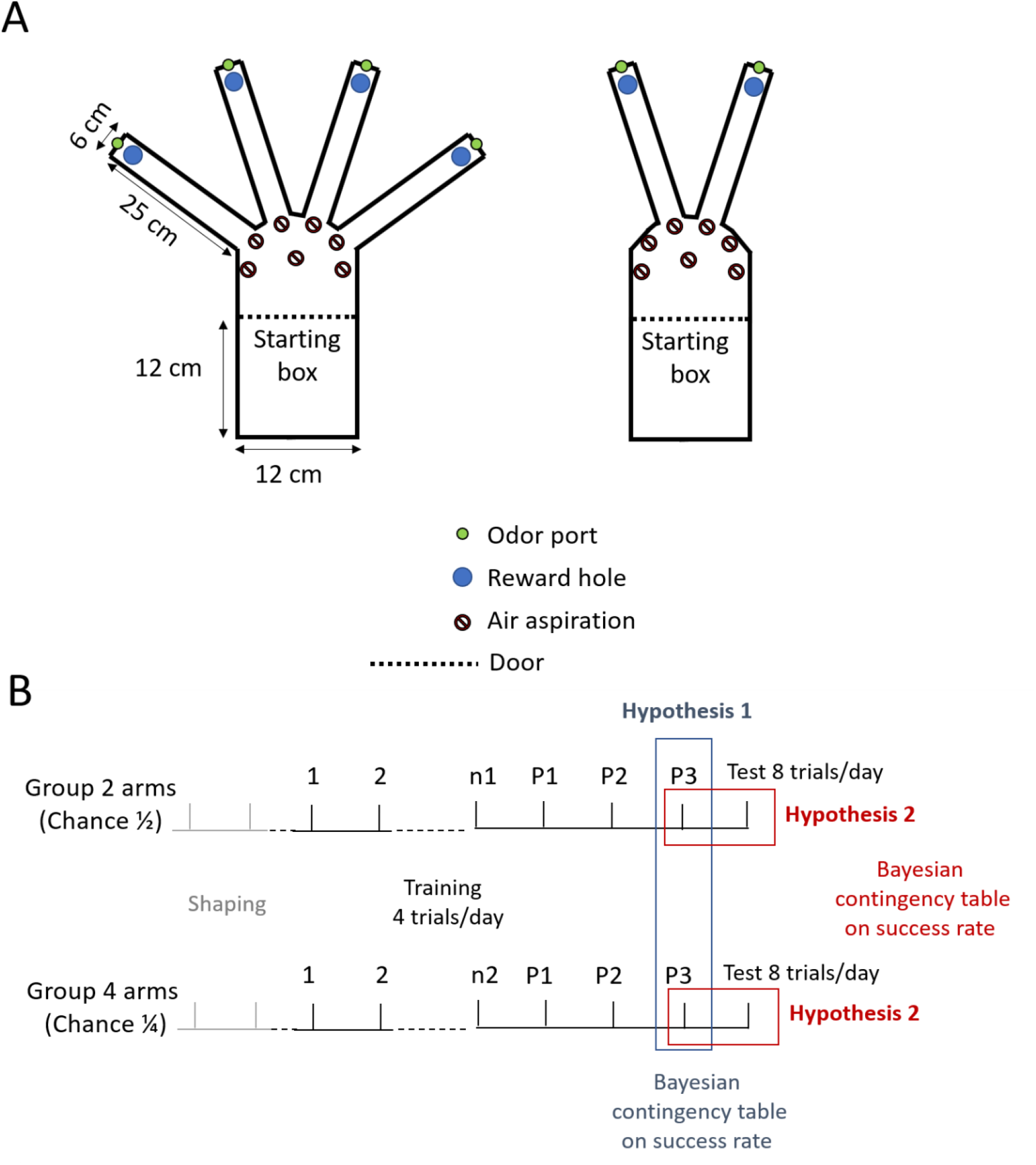
Experimental protocol. (**A)**, Radial mazes for the behavioral experiments. The holes are 8 cm deep with a diameter of 31 mm. Left, chance level of ¼. Right, chance level of ½. (**B)**, Behavioral planning involved statistical tests to verify the hypothesis. Tests were conducted once the success rate reached a plateau, defined as no modification of the success rate exceeding 5% over previous sessions for three consecutive sessions (P1-P3). A supplementary session with 8 trials was performed thereafter to test whether the animals’ success rate was modified by the increase in the number of trials. n1 and n2, last training day before plateau for group 1 and 2 respectively. P1-P3, training when success rate reaches the plateau.

During the subsequent training phase, mice were initially placed in a closed starting box for approximately 10 seconds. After this period, the door leading to the maze was opened, allowing the mice access. A food reward was randomly positioned at the end of one of the maze arms. This rewarded arm was scented with limonene through an odorant port that remained open throughout the trial. Mice remained in the maze until they located the reward, with a “successful” trial occurring when the arm containing the reward was the first one visited by the animal.

Two groups of 6 mice each were used. Group 1 underwent training in a two-arm maze (with a chance level of ½), while Group 2 trained in a four-arm maze (with a chance level of ¼). Each animal underwent one training session per day consisting of four trials per session with the animals from the two groups alternating. A trial was considered successful when the entire body of the animal entered the arm. For each session and each group, the same random sequence of rewarded arms was used, with this sequence eventually changing between sessions. A pseudo-randomization was employed to prevent the same arm from being rewarded more than twice consecutively in each session. Immediately after they had heated the reward, the animals were placed in a transfer cage while the maze was reset. This took a few tens of seconds. The entire maze was cleaned with alcohol after each animal completed the session. For each group, training continued until the average success rate reached a plateau (no modification of success rate > 5% over previous sessions for three consecutive sessions). After the third session at the plateau, a last session consisting of 8 trials was performed. Detailed information on the task and animals’ performance is available at osf.io/rt7d5/.

#### Tested hypothesis and statistic

Two experimental hypotheses were tested:

Hypothesis 1: The reduction of the chance level does not reduce the animal success rate. The alternative hypothesis, “success rate of group 1 > success rate of group 2”, was used to test the null hypothesis, “success rate of group 2 ≥ success rate group 1”, by using Bayesian contingency table (JASP Team, 2022). Bayes Factor (BF)01 >3 was the threshold to accept the null hypothesis (Keysers et al., 2020).

Hypothesis 2: The increase in the number of trials does not reduce the animal success rate. The alternative hypothesis,” success rate with 4 trials > success rate with 8 trials”, was used to test the null hypothesis, “success rate with 8 trials ≥ success with 4 trials”, by using Bayesian contingency table (JASP Team, 2022). BF01 >3 was the threshold to accept the null hypothesis. The default prior concentration of 1 was used for the Bayesian analysis

The absence of a reduction in success was also assessed using the frequentist TOST test (not pre-registered). For this we selected a low equivalence margin of 10% for the test, as even a reduction in the success rate of this magnitude—caused either by the modification of the chance level from 1/2 to 1/3 or by the increase in the number of trials from 4 to 8—still results in an increase in statistical power (see Figure 3E and https://osf.io/x6cey/ “Success rate” folder)

All JASP files related to the statistical analysis are available here.

#### Analytical procedure

For each session, the success rate of each animal is calculated as the sum of successes divided by the total number of trials. Descriptive statistics, including the average success rate of the population and its corresponding 95% confidence interval (calculated as the standard error of the mean multiplied by 1.96), are then presented for the different training sections. Statistical differences from the chance level were assessed using a one-tailed Student’s t-test.

## Results

In order to determine the statistical power for studies that aim to evaluate success rates, we have developed, a power calculator based on Monte Carlo simulation freely available here SuccessRatePower. Here a simple, notated interface allows users to define the different parameters of the experimental plan (Figure. 4A). We used SuccessRatePower to assess the impact of modifying the different parameters shown in Figure 1A on statistical power. Simulations were run under two conditions: in the first, the success rate of the experimental group was compared either to chance or to the performance of a second group performing at chance (Figure 4B). In the second, the comparison was made between two groups both performing above chance (Figure 5A).

**Figure 4:**
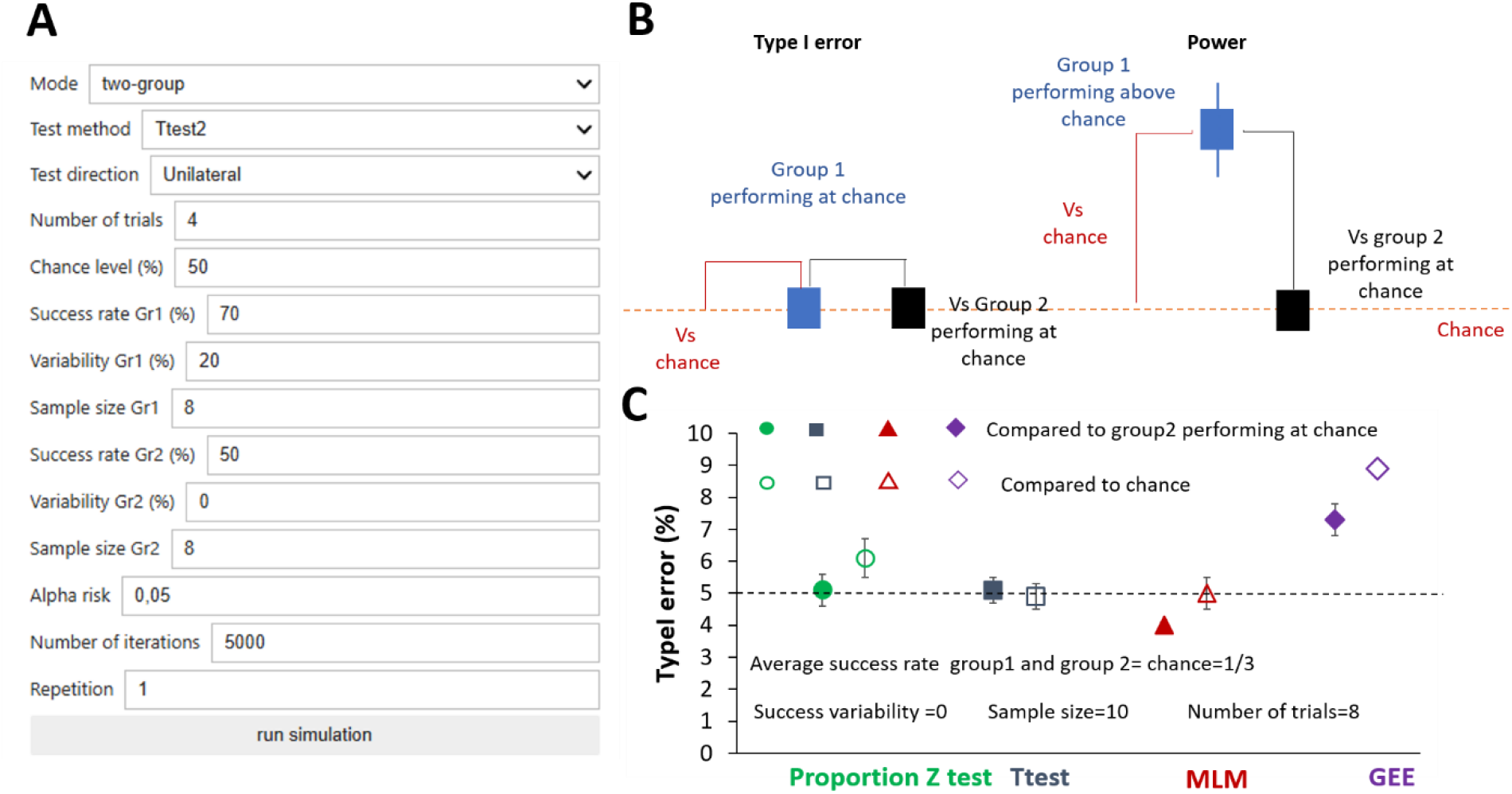
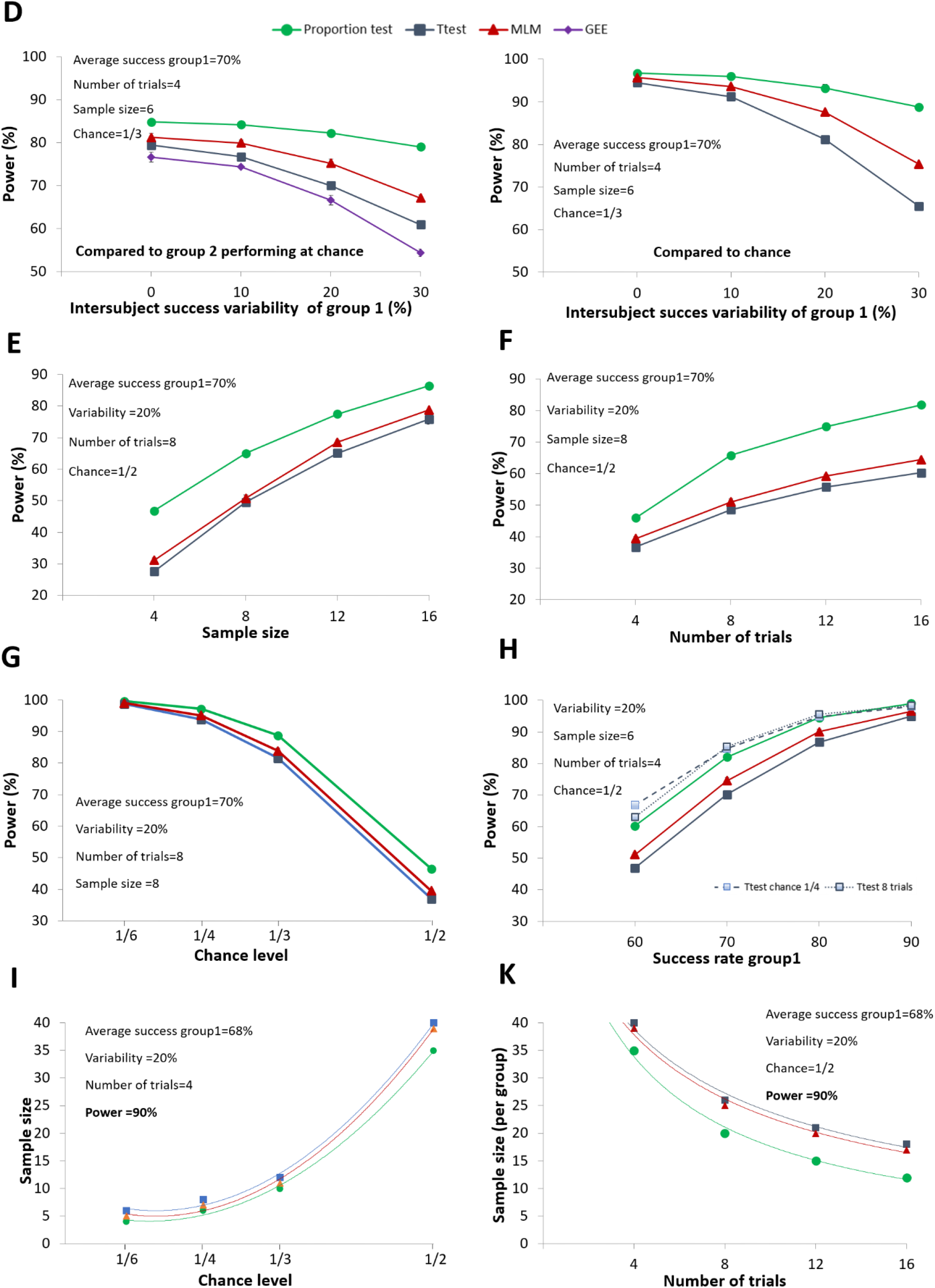
Type I error and statistical power as a function of different experimental parameters when one group is compared to chance or control group performing at chance level. **(A)** *SuccessRatePower* interface outlining the parameters employed in the simulation to compute statistical power (see Methods for more details). **(B)** Graphical representation of the experimental configuration used for Type I error and power calculation. **(C)** Occurrence of Type I error as a function of the analytical procedure. **(D)** Influence of inter-subject success variability on statistical power. Comparison between the two groups on the left, and between one group and the chance level on the right. **(E)** Statistical power evolution as a function of sample size. **(F)** Statistical power evolution as a function of number of trials. **(G)** Statistical power evolution as a function of chance level. **(H)** Statistical power evolution as a function of success rate and the modification of chance level or trial number (only for Ttest) **(I)** Evolution of the number of subjects required to obtain a statistical power of 90% as a function of the chance level. **(K)** Evolution of the number of subjects required to obtain a statistical power of 90% as a function of the number of trials. Notably, when not visible the 95% confidence interval bars are smaller than the size of the markers. The data corresponding to these figures can be accessed at (https://osf.io/gqybv). The script for the simulation can be found at (https://osf.io/ec7mp). MLM = Multilevel Mixed Model; GEE = Generalized Estimating Equations.

**Figure 5:**
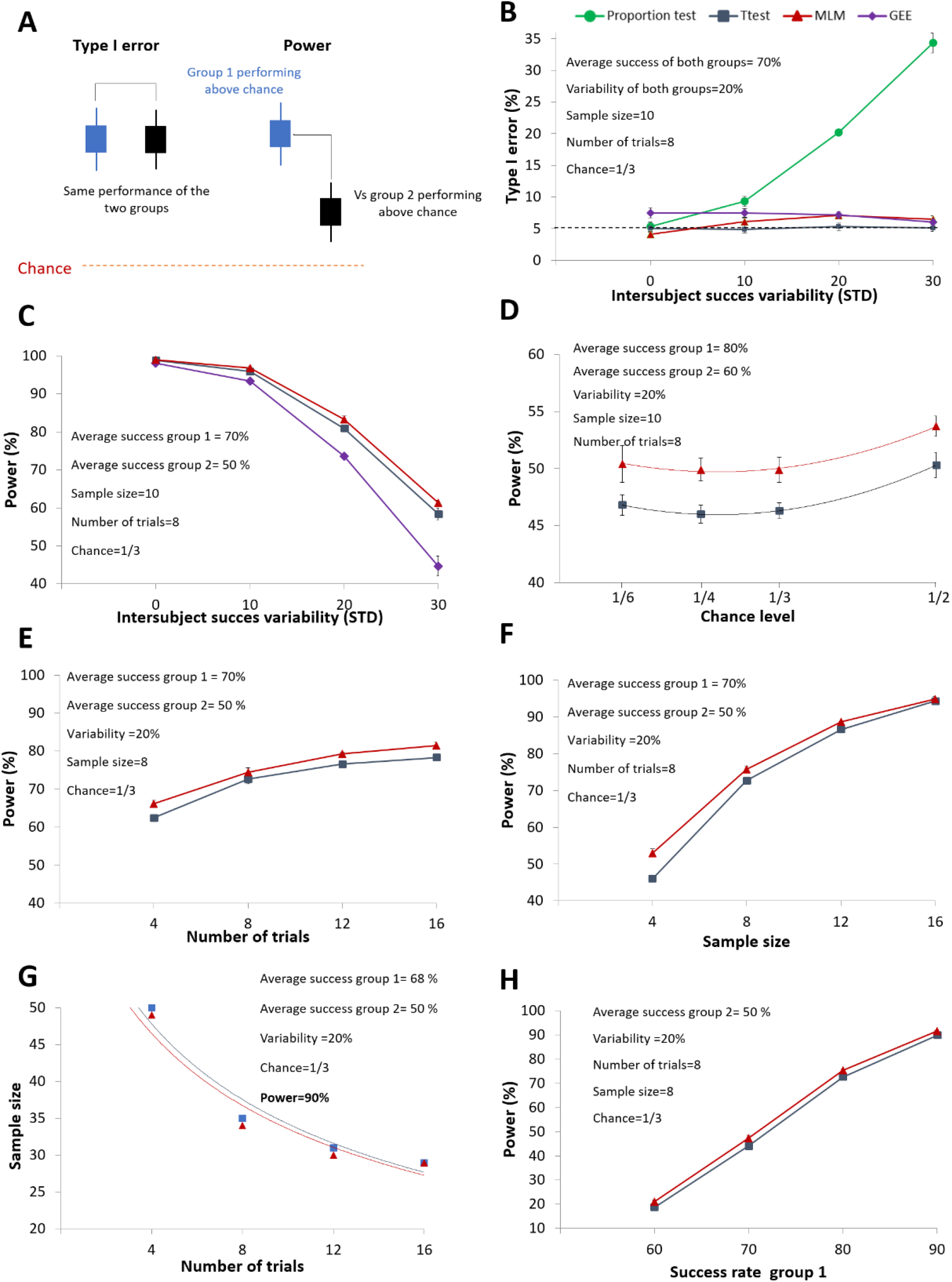
Type I error and statistical power as a function of different experimental parameters when one group is compared to control group performing above chance level. **(A)** Graphical representation of the experimental configuration used for Type I error and power calculation. **(B)** Occurrence of Type I error as a function of the analytical procedure. **(C)** Influence of inter-subject success variability on statistical power. **(D)** Statistical power evolution as a function of chance level. **(E)** Statistical power evolution as a function of number of trials. **(F)** Statistical power evolution as a function of sample size. **(G)** Evolution of the number of subjects required to obtain a statistical power of 90% as a function of the number of trials. **(H)** Statistical power evolution as a function of success rate. Notably, when not visible the 95% confidence interval bars are smaller than the size of the markers. The data corresponding to these figures can be accessed at (https://osf.io/gqybv). The script for the simulation can be found at (https://osf.io/vsuja/). MLM = Multilevel Mixed Model; GEE = Generalized Estimating Equations.

### Determinants of statistical power in experimental condition 1

Figure 4C illustrates the occurrence of Type I errors under the first condition. In this simulation, the success rates of both the treated and control groups were set at chance level, with inter-subject variability fixed at zero. When two groups performing at chance were compared, the Type I error rate remained at the nominal α level when using either proportion-based tests or a two-sample t-test. With a multilevel mixed-effects model, the error rate was slightly below α, whereas a small inflation was observed with generalized estimating equations (GEE) (Figure 4C, filled markers). Conversely, when the experimental group was compared directly to chance, the Type I error was slightly inflated with the proportion test and more markedly with GEE (Figure 4C, empty markers).

Figure 4B show the decrease in statistical power as the variability of success rates between subjects in the treated group increases. As expected, statistical power was higher when comparing the success rate of the experimental group directly to the chance level (right panel), rather than comparing it to a control group performing at chance level (left panel). This difference arises because statistical power declines with greater data variability, which in turn increases when data come from two groups. For the remaining simulations, only comparisons with a control group will be illustrated. Moreover, due to its high Type I error rate and low statistical power, the performance of the GEE model was not further investigated. Overall, among the analytical methods tested, the procedure based on the proportion test provided the highest statistical power, particularly when inter-subject variability in success rates increased. Figures 4E and 4F show that statistical power increases with larger sample sizes and greater numbers of trials, respectively. Notably, increasing the sample size yields a stronger improvement in power than increasing the number of trials, although this difference is less pronounced when the proportion test is applied. The increase in statistical power with a greater number of trials is attributable to the improved estimation of the true success rate. As a result, the differences between subjects’ success rates reflect more of the true within-subject variability, rather than being inflated by experimental noise due to too few trials (Supplementary Figure 4). As shown in Figure 4G the statistical power strongly increases with the decrease of the chance level while Figure 4H illustrates how statistical power changes with the average success rate of the treated group. Notably, modifying the chance level or the number of trials has little impact on statistical power when the statistical power is equal to or greater than 90%.

Overall, when the experimental group is compared to chance or when the control group is expected to perform at chance level, statistical power can be increased by reducing the chance level and by increasing the number of trials performed by the subjects without the need to increase the sample size. Figures 4I and 4K respectively show the reduction in the number of subjects required to reach 90% power when the chance level decreases or the number of trials increases. Concerning the analytical methods, the use of the proportion test shows better performance in terms of statistical power, especially when inter-subject variability and/or the number of trials is high. However, it produces a slight inflation of the Type I error rate when the treated group is compared to chance. In such cases, the use of multilevel linear models or analytical procedures designed for continuous data (e.g., the t-test) should be preferred.

### Determinants of statistical power in experimental condition 2

When both experimental groups perform above chance (Figure 5A), the use of the analytical procedure based on the proportion test leads to a consistent inflation of the Type I error rate, which increases with greater variability in success rates across subjects. In contrast, for the other three methods, the Type I error rate remains close to the nominal α level—exactly at α for the t-test, and slightly above it for the MLM and GEE approaches (Figure 5B). Figure 5C shows that statistical power decreases as inter-subject variability increases. Notably, the GEE approach performs significantly worse than the t-test and MLM in this context. Because of inflated Type I error rates or poor statistical power, the analytical approaches based on the proportion test and GEE were not considered further. Interestingly, in this experimental condition, modifying the chance level no longer influences statistical power. Moreover, the MLM approach shows a slight increase in power compared to the t-test method (Figure 5D). Additionally, increasing the number of trials continues to improve statistical power (Figure 5E). Although this gain is smaller than the one achieved by increasing the sample size (Figure 5F), running experiments with more trials can substantially reduce the number of subjects needed to achieve high statistical power (Figure 5G). Finally, Figure 5H illustrates how statistical power changes as a function of the difference in average success rate between the two groups.

In conclusion, when both groups perform above chance, increasing the number of trials enhances statistical power, whereas modifying the chance level does not. In addition, proportion-based analyses should be avoided to prevent inflated Type I error rates.

In summary, these simulations demonstrate that designing behavioral studies with a minimal chance level or a high number of trials, can lead to achieving a high level of statistical power, even with a low sample size. However, this conclusion is contingent upon ensuring that these parameters do not compromise subject performance since, as illustrated in Figure 4H and 5H, statistical power decreases with a reduction in subject success rate. Indeed, we cannot a priori exclude the possibility that a decrease in the chance level and/or an increase in the number of trials may impact subject performance. For instance, lowering the chance level may heighten task difficulty, consequently reducing subject success rates. Additionally, increasing the number of trials may lead to decreased success rates due to factors such as fatigue or diminished motivation (e.g., in animal studies utilizing food rewards where satiety can affect subject engagement). Therefore, to experimentally test these eventualities, we conducted an associative task in mice by varying these two parameters.

### Animal performance is not reduced by decreasing the chance level or increasing the number of trials

To assess the impact of modifying the chance level on animal performance, two groups of adult mice (six animals per group) participated in an associative olfactory task within a radial labyrinth, where they encountered either two arms (chance ½) or four arms (chance ¼). The mice underwent training until performance plateaued. The absence of differences in success rates between the two groups was evaluated using a Bayesian statistical test, calculating the probability of the null hypothesis over the alternative (BF01). The training session consisted of four trials, except for the last session where eight trials were used in order to evaluate the impact of altering the number of trials on success rates, (for further details, refer to the methods section and Figure 3).

Both the behavioral protocol and analytical plan were pre-registered (here, OSF). Figure 6A illustrates that both groups displayed similar learning curves, with the ¼ chance level group reaching statistically significant effects earlier than the ½ chance level group. Additionally, both groups exhibited comparable performance at the plateau (87.5± 14% for ¼ chance level group, 87.5 ± 20% for ½ chance level group at training session 15; BF01=4.2, p=0.15 TOST test, n=6). Moreover, the success rate remained unaffected by doubling the number of trials performed by the mice during the eight-trial session (For two arms group: 87.5± 20% for 4 trials, 91.7 ± 10% for 8 trials; BF01=8.7, p=0.02 TOST test. For four arms group: 87.5± 14% for 4 trials 87.5± 14% for 8 trials BF01=5.3, p=0.06 TOST test, n=6).

**Figure 6:**
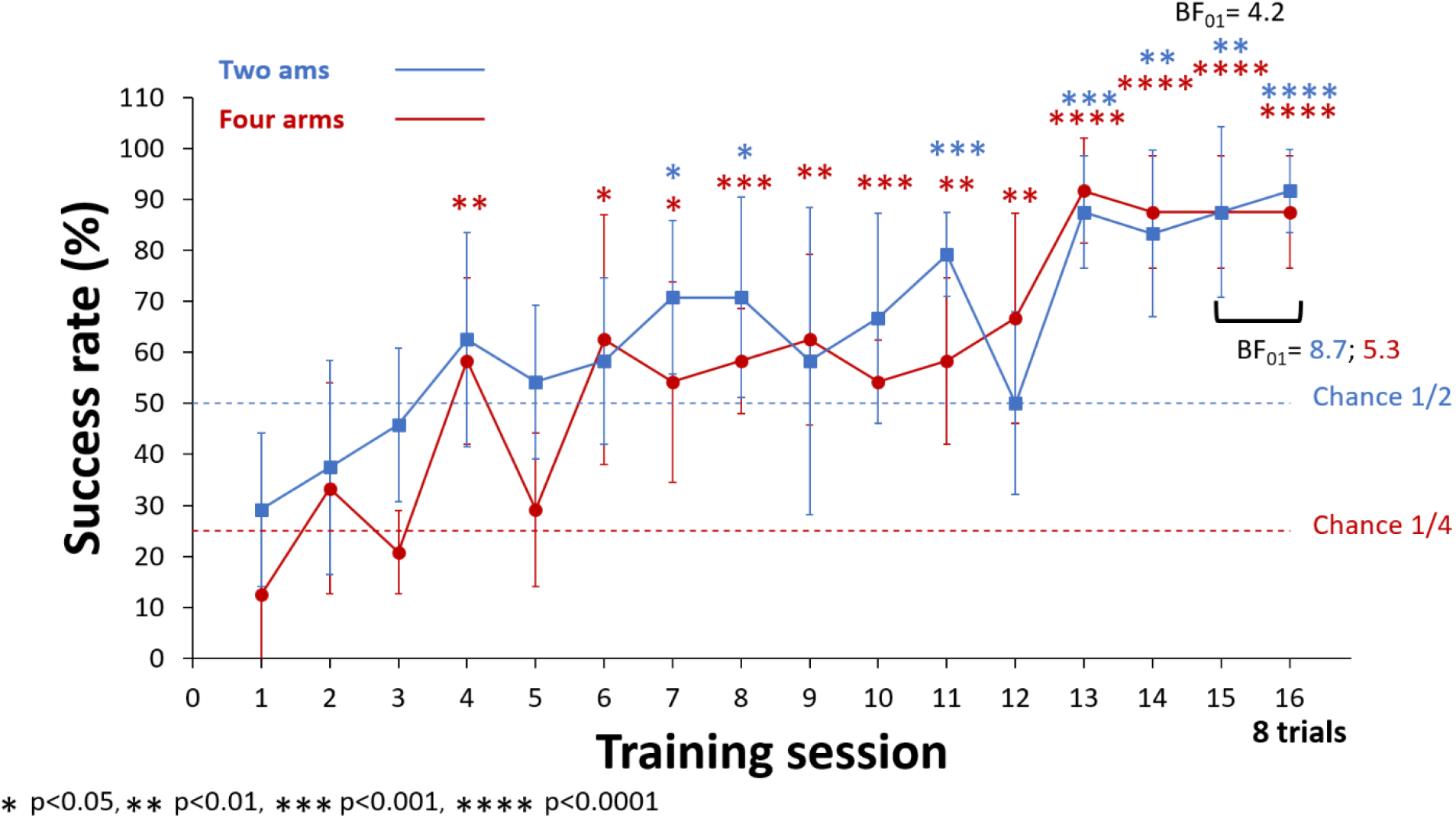
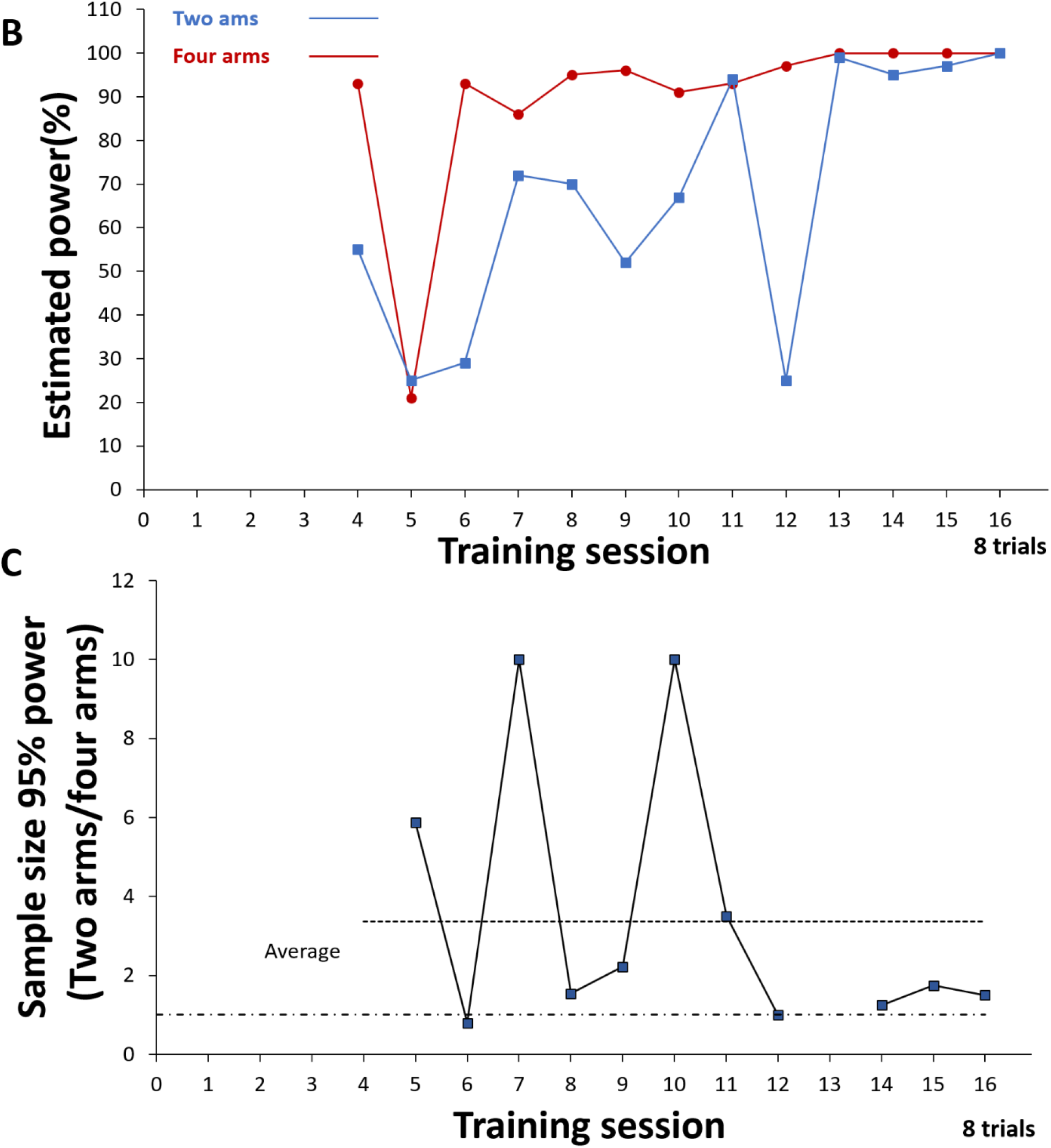
Success rate is not affected by changing the chance level or the number of trials performed by the mice. (**A)** Learning curve of mice performing in a two-arm maze (chance level ½) or a four-arm maze (chance level ¼). Mice underwent four trials per day until reaching session 16. Asterisks denote statistical significance compared to chance. The Bayesian factor indicates the absence of a significant difference between the two groups at training session 15, as well as the absence of a difference between four and eight trials within each group. **(B)** Estimated statistical power calculated with the data presented in (A). **(C)** Estimation of the increase in animal requirements to reach 95% statistical power when the task is performed in a two-arm (chance level ½) versus a four-arm (chance level ¼) condition. Lower dashed line depict ratio =1. All data and statistical analysis are available at https://osf.io/rt7d5/.

Overall, these results suggest that, at least for the tested conditions, the success rate of the experimental groups is not modified either by the reduction of the chance level or by the increase in the number of trials, suggesting that the modification of these two parameters is a good strategy to increase statistical power.

The implementation of SuccessRatePower enabled the utilization of experimental data to estimate statistical power across the session. As depicted in Figure 6B, the estimated power is clearly larger for the four-arms design. This superiority is particularly evident in session with lower average success rates. Additionally, we utilized SuccessRatePower to estimate the necessary number of subjects for achieving 95% power, and to assess the ratio of animal requirements between the four-arms and two-arms conditions (Figure 6C). This analysis shows that, on average, three times more animals would be necessary when the task is performed with chance level is ½ compared to the condition where task is performed with chance ¼.

## Discussion

In this report, we present ‘SuccessRatePower’ a new statistical tool designed to calculate statistical power in studies evaluating population success rates, such as those conducted in behavioral neuroscience assessing differences in cognitive abilities. This software considers various parameters of the experimental design, allowing researchers to evaluate the impact of their modifications on power and thus select the optimal experimental configuration for these studies. Using this tool, we demonstrated that, beyond increasing the sample size, statistical power can also be improved by adjusting three parameters under the experimenter’s control: (1) increasing the number of trials used to estimate success rates, (2) lowering the chance level—particularly when the control group is expected to perform at chance—and (3) to a lesser extent, applying alternative statistical analyses better suited to the nature of the experimental outcomes. In this regard, it is noteworthy that the majority of published studies are designed using the higher chance level (½), even when adjusting this parameter can significantly influence statistical power. Additionally, statistical analyses are often performed after transforming discrete data into continuous variables. These practices may stem from historical conventions and methodological traditions, which together may have contributed to a lack of awareness regarding the impact such transformations have on statistical power.

The prevalence of underpowered design in the literature may also be attributable to the anecdotally common concern that decreasing the chance level or increasing the number of trials could compromise the subject’s performance and reduce its success rate. Here, we provide experimental evidence demonstrating that, at least for simple associative tasks, this is not the case.

Therefore, by simply modifying a few parameters within the experimental design, we can significantly reduce the number of animals used without compromising the quality of the statistical inference. For example, in the study by Mandairon et al. (2018), the associative task comparing the performance between trained and pseudo-trained animal was designed with 4 training trials with a chance level of ½. After 5 days of training, the success rates for the trained and control groups were 77% and 49%, with success variability at std=19 % and std=20 %, respectively. Using 20 animals per group, the statistical power achieved was 92%. However, “SuccessRatePower” demonstrated that the same power could have been reached with only 5 mice per group by performing the experiment with 8 trials at a chance level of ¼. This adjustment not only saves time, but also provides a significant ethical benefit, as minimizing the number of animals necessary for a given protocol is a cornerstone of ethical scientific practice.

It should be noted that the advantages produced by modifying the aforementioned parameters diminish as the success rate of the trained group increases, becoming negligible for values greater than 90% (Figure 4H). Moreover, the benefit of using statistical methods for discrete values decreases as the variability in performance between subjects is reduced (Figure 4D). Importantly, that statistical analyses for discrete values can only be performed if all subjects undergo the same number of trials, to avoid results being driven by the performance of over-tested subjects.

The interventions in the experimental design that can lead to an increase in statistical power in behavioral experiments evaluating success rates are resumed in table 2.

**Table 2:** Possible interventions in the experimental design that lead to an increase in statistical power without the need to increase the sample size.

| INTERVENTION | EXPERIMENTAL CONDITION | CAUTIONARY POINTS |
| --- | --- | --- |
| *Increase of the number of trials* | All conditions | Verify the possible reduction in subjects' performance and its impact on statistical power. |
| *Use of lower chance level* | When the experimental group is compared to chance.<br>When the control is expected to perform at chance level. | Verify the possible reduction in subjects' performance and its impact on statistical power |
| *Use of statistical test suited for proportion comparison* | Only when one of the groups is expected to perform at chance level | Each subject must perform the same number of trials |

Overall, in these types of studies, the parameters influencing statistical power are numerous and interdependent. Conventional power calculators based on mathematical approaches can account for only a limited set of specific conditions. In contrast, the SuccessRatePower software, which leverages Monte Carlo simulations, addresses these limitations by allowing users to evaluate the impact of modifying multiple design parameters on statistical power.

It is important to highlight, however, that SuccessRatePower has three limitations compared to conventional power calculators:

1. Its use is restricted to behavioral experiments with the design outlined in Figure 1.

2. It provides an approximation of the exact statistical power, albeit with minimal variability when the number of iterations is set to 5,000 or more (Supplementary Figure 2).

3. It cannot directly calculate the sample size required to achieve a specific statistical power. Users can, however, determine this by running simulations multiple times with different sample sizes.

Thus, for specific experiments evaluating and comparing subject success rates, the “SuccessRatePower” software is a valuable tool for optimizing the experimental protocol design.

## Acknowledgements

We thank Thomas Orset, Clémence Long, and Lucie Plantade who performed preliminary experiments on the behavioral task

## Funding

The authors declare that they have received no specific funding for this study.

## Conflict of interest disclosure

The authors declare of having no financial or non-financial conflicts of interest in relation to the content of the article.

## Data, scripts, code, and supplementary information availability

Scripts are available [here](https://github.com/crnl-lab/StatistcalPowerCalculation_2024_NicolaKuczewski), row data and analysis [here](https://osf.io/sbm9e/) and supplementary figures [here](https://osf.io/wjzkq).

## Notes

https://osf.io/sbm9e/

### Competing Interest Statement

The authors have declared no competing interest.

### Summary of Updates

Addition of PCI Neuroscience badge and removal of line numbers

