## Supplementary figures for "Enhancing Statistical Power While Maintaining Small Sample Sizes in Behavioral Neuroscience Experiments Evaluating Success Rates" (from another version of this preprint: version 5, 2025-06-16)

A

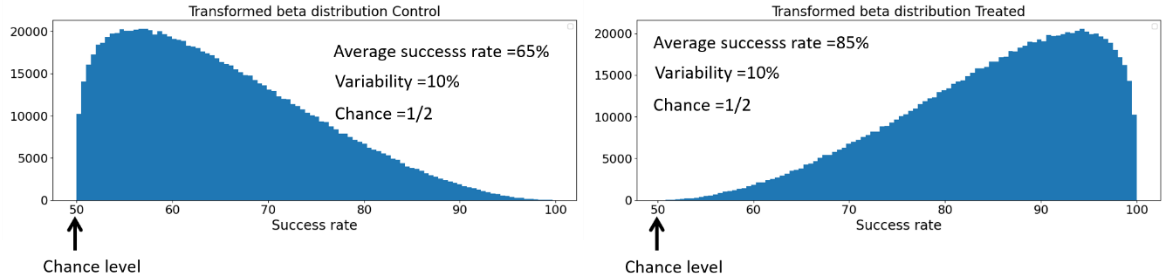


B

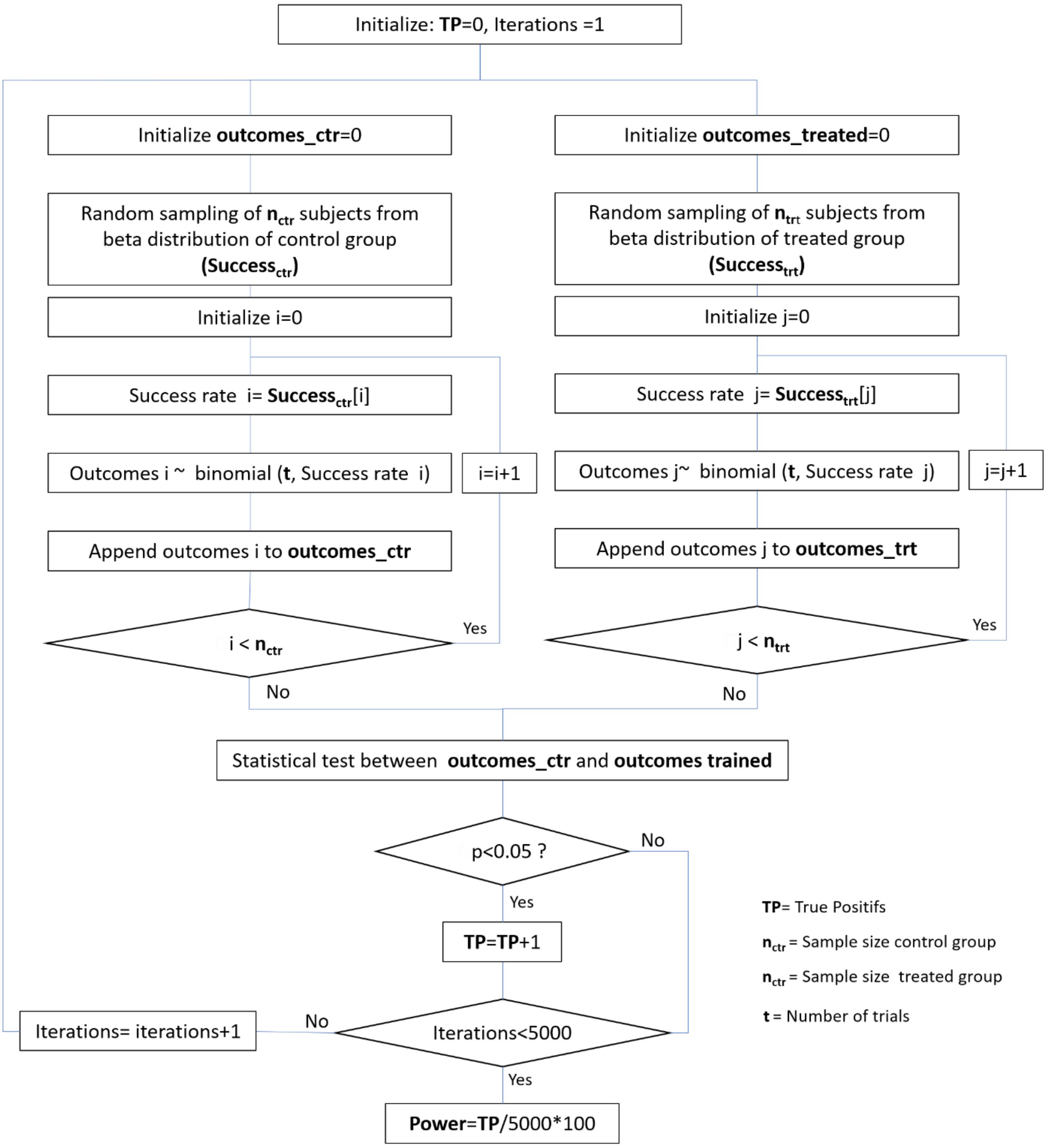


**TP**= True Positifs

$n_{ctr}$ = Sample size control group

$n_{ctr}$ = Sample size treated group

**t** = Number of trials

**Supplementary Figure 1:** Procedure for determining statistical power through Monte Carlo simulation **(A)** Example of transformed beta distributions used for random sampling of success rates. Noted that success rate cannot be lower than chance or higher than 100% **(B)** Flowchart of the Monte Carlo simulation for power calculation when the treated group is compared to a control group with a success rate above chance (by use of 5000 iterations). When comparing the treated group to chance, the control group is replaced by the chance-level value. When comparing to a control group performing at chance level, the success of each subject in the control group is set to the chance level. When both groups exhibit the same success rate, the script computes the occurrence of Type I errors (i.e., true positives become false negatives).

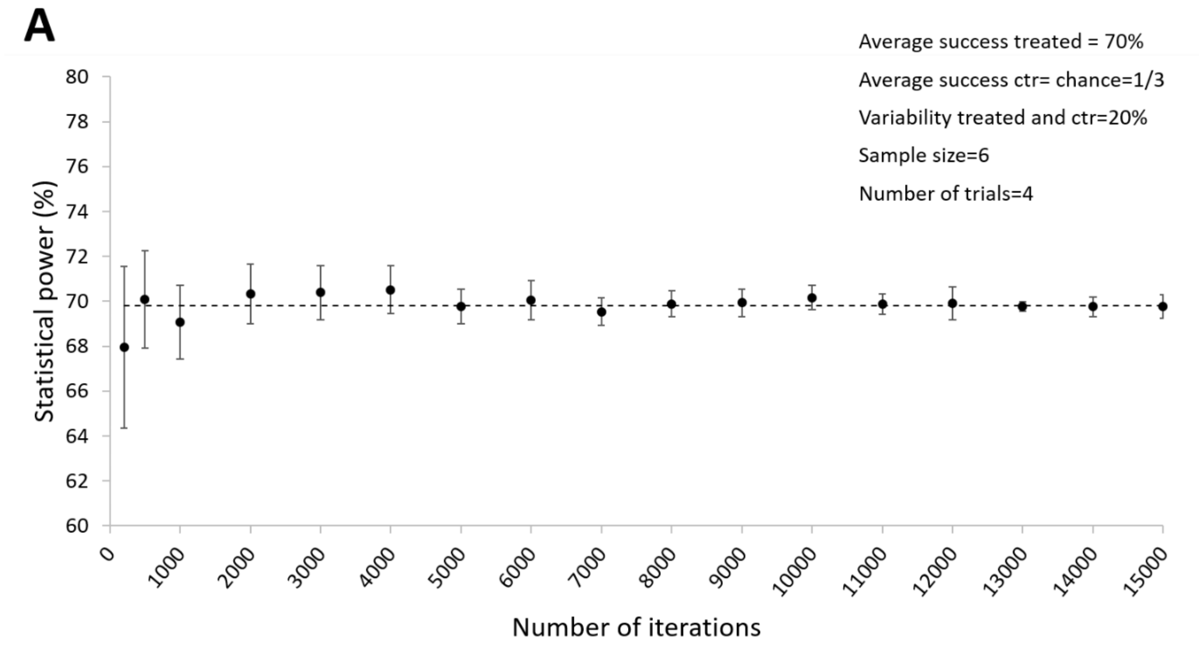


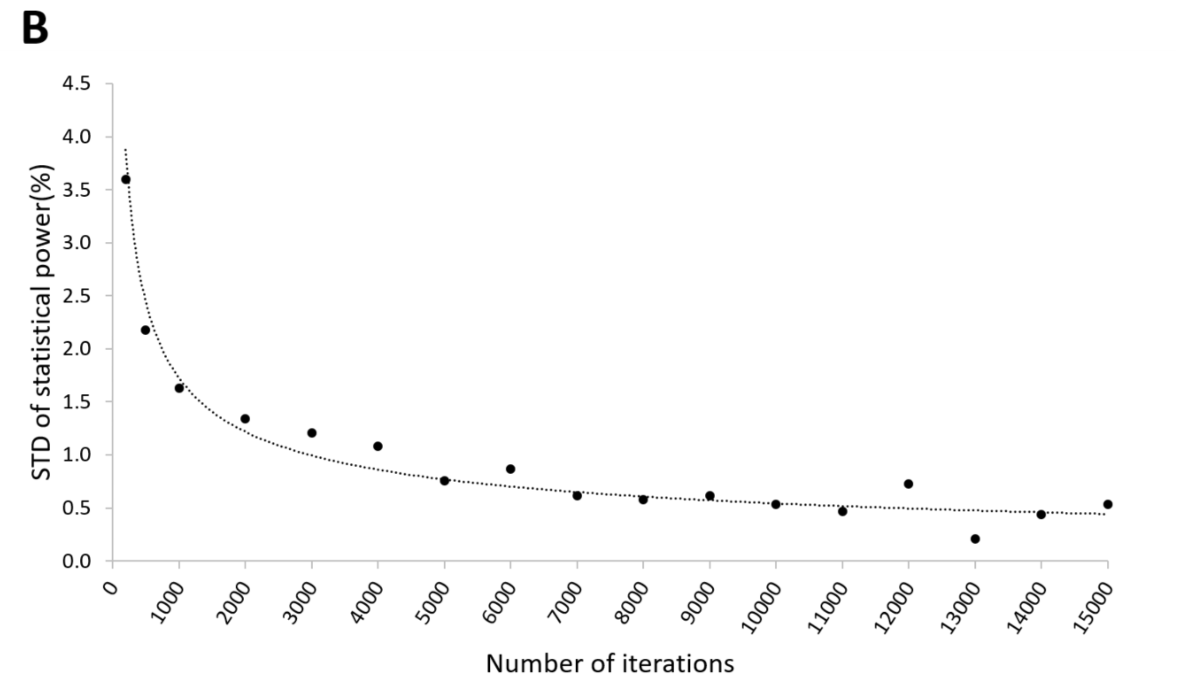

**Supplementary Figure 2**: Effect of the number of SuccessRatePower iterations on the variability of power estimation. **A)** Statistical power and its standard deviation as a function of the number of iterations used in the Monte Carlo simulation. Each point represents the average of 10 repetitions of power calculations (using the Ttest for comparing two groups). **B)** Evolution of the variability in power estimation as a function of the number of iterations used in the Monte Carlo simulation. Each point represents the standard deviation values shown in panel A. Dashed line in represent the average power in A and the exponential interpolation of the data in B.

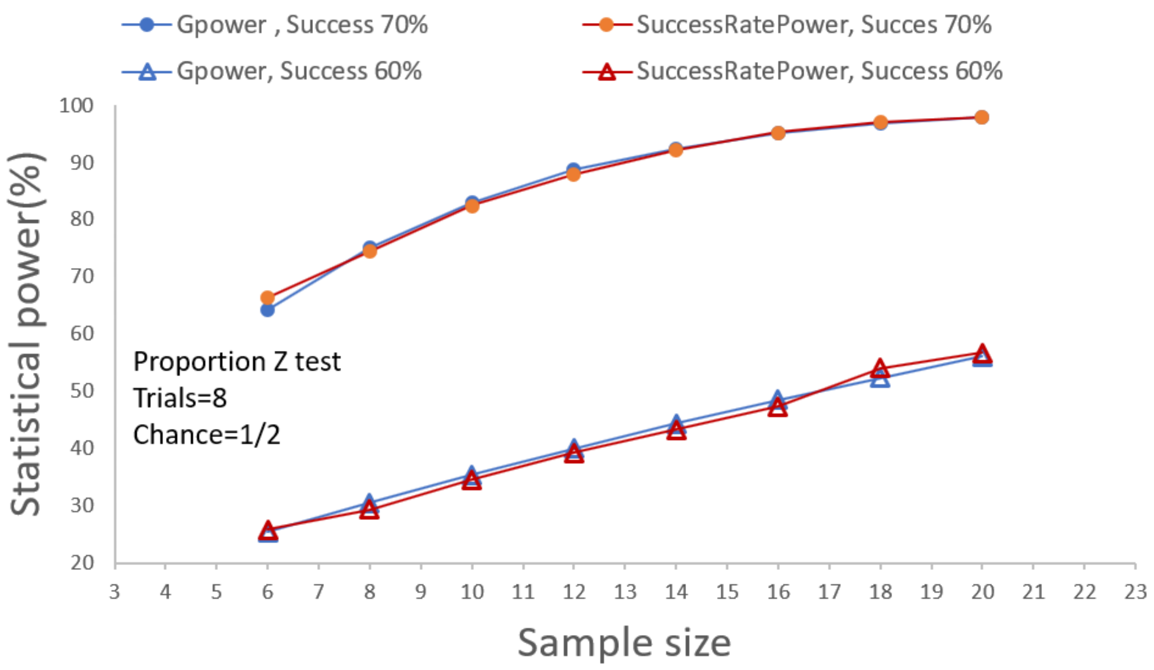


**Supplementary Figure 3** SuccessRatePower demonstrates similar performance to G*Power 3 under the specific experimental and analytical conditions where power calculation is feasible with G*Power 3 namely a proportion z-test comparing two groups with no intrasubject variability.

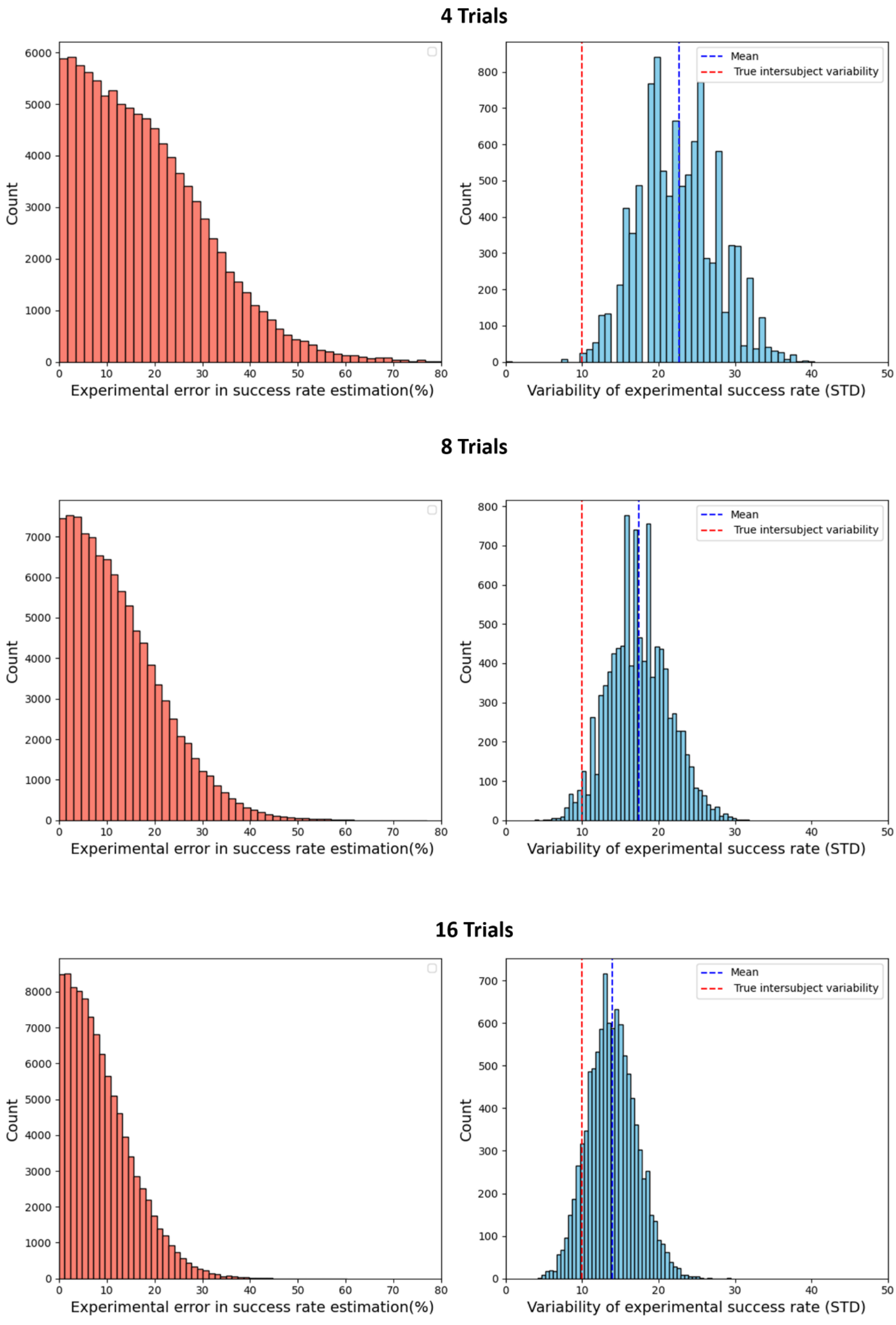


**Supplementary Figure 4**: As the number of trials in a session increases, the estimation of each subject's success rate becomes more accurate (left). This reduces the apparent variability between subjects, making it more reflective of their true intrasubject variability (right), leading to the increase of statistical power. (Success rate =70%, variability=10%, n=10, chance=1/3)
